# The reference genome for the northeastern Pacific bull kelp, *Nereocystis luetkeana*

**DOI:** 10.1101/2025.07.16.665147

**Authors:** Cicero Alves-Lima, Gabriel Montecinos, Merly Escalona, Sara Calhoun, Mohan Marimuthu, Oanh Nguyen, Eric Beraut, Anna Lipzen, Igor V. Grigoriev, Peter Raimondi, Sergey Nuzhdin, Filipe Alberto

## Abstract

Bull kelp, *Nereocystis luetkeana,* is a northeastern Pacific kelp with broad distribution from Alaska to central California. Its population declines have caused severe concerns in northern California, the Salish Sea in Washington, and recently in some populations in Oregon. Despite bull kelp’s accumulated ecological and physiological studies, an assembled and annotated genomic reference was still unavailable. Here, we report the complete and annotated genome of *Nereocystis luetkeana*, produced by the California Conservation Genomics Project (CCGP), which aims to reveal genomic diversity patterns across California by sequencing the complete genomes of approximately 150 carefully selected species. The genome was assembled into 1,562 scaffolds with 449.82 Mb, 80x of coverage and 22,952 gene models. BUSCO assembly showed a completeness score of 72% for the stramenopiles gene set. The mitochondria and chloroplast genome sequences have 37 Mb and 131 Mb, respectively. The orthology analysis between 10 Phaeophycean genomes showed 1,065 expanded and 286 unique orthogroups for this species. Pairwise comparisons showed 542 orthogroups present only in *N. luetkeana* and *M. pyrifera*, another large-body kelp. The enrichment analysis of these orthogroups showed important functions related to central metabolism and signaling due to ATPases enrichment in these two species. This genome assembly will provide an essential resource for the ecology, evolution, conservation, and breeding of bull kelp.

## INTRODUCTION

*Nereocystis luetkeana* is a canopy-forming large phaeophyceae found from Point Conception in California to Umnak Island, Alaska (Druehl 1970; Miller and Estes 1989; Konar et al. 2017; Pfister et al. 2018). Despite its wide distribution, recent population declines have caused serious concern in northern California (Rogers-Bennett and Catton 2019), the Salish Sea in the northeast Pacific (Berry et al. 2021), and some populations in Oregon (Hamilton et al. 2020). Marine heat waves caused by anthropogenic climate change originated a cascade of events that are associated with this decline (Assis et al. 2018; Wernberg et al. 2018; Smale 2020; Finger et al. 2021; McPherson et al. 2021). However, as a ruderal species, *N. luetkeana* can rapidly replenish dense beds even on resilient urchin barrens (Breen et al. 1976; Foreman 1977; Pace 1981; Dayton et al. 1984; Dayton 1985) if temperatures are optimal (Lüning and Freshwater 1988; Supratya et al. 2020).

Bull kelp sporophyte can grow up to 27.1 cm daily, reaching ten times its length and weight in less than two months (Foreman 1970; Kain 1987; Maxell and Miller 1996) and a total length of 40 meters or more (Setchell 1908). Among all known life forms, this is one of the highest mass-specific growth rates (Lynch et al. 2022). Morphological and physiological studies on bull kelp development have been described in detail (Nicholson 1970; Schmitz and Srivastava 1976; Duncan and Foreman 1980; Kain and Norton 1987; Koehl et al. 2008; Knoblauch et al. 2016b, a; Liggan and Martone 2020), as much as its high nutrient uptake rates (Wheeler et al. 1984; Ahn et al. 1998), and tolerance to salinity (Lind and Konar 2017), mechanical (Koehl and Wainwright 1977; Koehl and Alberte 1988; Denny et al. 1997) and light (Popovic et al. 1983; Poulson et al. 2011) stresses.

As an annual species, *N. luetkeana* is known for its rhythmic phenology, showing diel spore release (Amsler and Neushul 1989) and tidal sori abscission (Walker 1980) that is expected to produce a bimodal spore release height (Amsler and Neushul 1989; Burnett et al. 2024). This sorus abscission-spore release mechanism was proposed to be an adaptation to maximize the photosynthetic potential of the spores and a dispersal life history where a proportion of the spores can travel long distances while others are retained near the parents (Amsler and Neushul 1989, Burnett et al. 2024). Population genetics analysis of the species across its range confirmed this life history (Gierke et al. 2023). This study revealed significant inbreeding coefficients, likely caused by the local retention of spores, while also showing decreased genetic differentiation across long stretches of coast.

Four main groups of genetic co-ancestry, genetic diversity hotspots, and impoverished areas, have been detected in bull kelp using microsatellite markers (Gierke et al. 2023). Nevertheless, characterizing genetic variants mapped to a reference genome can reveal additional substructuring (Bemmels et al. 2025) and, most importantly, allow for the detection of localized adaptation. This level of understanding is essential to guide which source should be used when restoring natural populations from biobanks of *ex-situ* preserved gametophytes (Wade et al. 2020).

Here we report the complete genome of *Nereocystis luetkeana*, assembled by the California Conservation Genomics Project (CCGP), which aims to reveal patterns of genomic diversity across California by sequencing the complete genomes of approximately 150 carefully selected species (Shaffer et al. 2022). This genome assembly will provide an essential resource for understanding bull kelp ecology, evolution, and conservation.

## METHODS

### Biological Materials

Fertile fronds of *Nereocystis luetkeana* (K. Mertens) Postels and Ruprecht (Lessoniaceae, Laminariales) sporophyte were sampled from North Beach, Port Townsend, WA (48.14533, –122.77633) in July 2019, and sent to the University of Wisconsin-Milwaukee where spore release was induced as previously described (Redmond et al. 2014). Developed female gametophytes were distinguished from males based on morphology, isolated, and submitted to the germplasm bank in UW-Milwaukee under code NB12FB3. Vegetative propagation was performed in 25 cm^2^ polystyrene vented culture flasks (Nunc) filled with 20 ml of sterile Provasoli media-enriched semi-artificial seawater (Instant Ocean Salt), with salinity 34, pH 8.2, and under 40 μmol photons·m^-2^·s^-1^ of irradiance from white fluorescent full spectrum bulbs covered with red cellophane paper. Bacterial contamination was decreased by treating with an antibiotics mix (ampicillin 125 μg·ml^-1^, ciprofloxacin 15 μg·ml^-1^, kanamycin 30 μg·ml^-1^, and streptomycin 30 μg·ml^-1^) for one week. The biomass was fragmented weekly with a sterile pestle to enhance vegetative growth up to 100 mg. At this stage, the tissue was freeze-dried and shipped in dry ice to UC Davis and UC Santa Cruz for HiFi and Omni-C library prep and sequencing respectively.

### Nucleic acid extraction

We extracted high molecular weight (HMW) genomic DNA from 693 mg of the tissue-cultured female haploid gametophyte using the cetyltrimethylammonium bromide (CTAB) method as described previously (Inglis et al. 2018), with the following modifications: (i) we used sodium metabisulfite (1% w/v) instead of 2-mercaptoethanol (1% v/v) in the sorbitol wash buffer and CTAB solutions; and (ii) repeated the tissue homogenate wash steps until the supernatant turned clear. We also performed (iii) the CTAB lysis at 45°C and (iv) the chloroform extraction twice using ice-cold chloroform. The extracted HMW DNA was purified using the high-salt-phenol-chloroform and phenol-chloroform methods (Pacific BioSciences – PacBio, Menlo Park, CA). The DNA purity was estimated by absorbance ratios (260/280 = 1.84 and 260/230 = 2.16) measured using the NanoDrop ND-1000 spectrophotometer (Thermo Fisher Scientific, Waltham, MA). The DNA yield (2.3 μg) was quantified using a Quantus Fluorometer (QuantiFluor ONE dsDNA Dye assay; Promega, Madison, WI), and the size distribution of the DNA was estimated using the Femto Pulse system (Genomic DNA 165 kb kit, Agilent, Santa Clara, CA), where 80% of the DNA fragments were found to be 30 kb or longer.

To assist the annotation of the bull kelp genome, we extracted RNA from male and female gametophyte tissue preserved in a biobank (samples NB12FB3 and NB12MB4). The gametophytes were exposed to red and white light for one week and sampled in the first hour of the day and the first hour of the night. The RNA was extracted from 20 mg of the tissue-cultured female haploid gametophyte sampled from T75 (20 ml) flasks with wide-bore 1 ml tips. Before weighing, the tuft moisture was removed by centrifugation for 2 min at 14,000 RPM and 4°C using silica spin columns (Macherey-Nagel). The tufts were removed with tweezers and weighed on an analytical scale. The tissue was transferred to 2 ml U-bottom tubes with two tungsten carbide beads (5 mm) and flash-frozen in liquid nitrogen. The tissue was ground thrice in frozen 24-tube racks of QIAGEN tissue-lyser for 1 min at 25 Hz. The racks were cooled in dry ice between each round for 5 minutes. The fine powder was diluted in the extraction buffer from the Macherey-Nagel Plant and Fungi (PN: 740120.50) extraction kit and incubated at 25°C for 40 min vortexing at 1,400 RPM. The following steps followed the manufacturer’s recommendations. NanoDrop spectrophotometry, QUBIT fluorimetry (RNA broad range kit), and 1.5% agarose gel electrophoresis were done to check RNA quality and integrity. All samples had A260/A280 and A260/A230 higher than 1.8 and a yield higher than 1 ug. The RNA was sent to the Gene Expression Center at the University of Wisconsin Biotechnology Center in Madison – WI.

### Omni-C library preparation

For chromosome conformation, an Omni-C library was prepared using the Dovetail™ Omni-C™ Kit (Dovetail Genomics, Scotts Valley, CA), following the manufacturer’s protocol with minor adjustments. First, specimen tissue (ID: NB12F.D) was thoroughly ground with a mortar and pestle while cooled with liquid nitrogen. Subsequently, chromatin was fixed in the nuclei to maintain its structural integrity during extraction, and then the chromatin solution was filtered through 100 μm and 40 μm cell strainers to remove debris. DNase I digestion was conducted under varied conditions for optimal DNA fragment size distribution. Chromatin ends were repaired and ligated to a biotinylated bridge adapter to facilitate proximity ligation. Following ligation, crosslinks were reversed, and DNA was purified to remove proteins. Biotin on non-ligated DNA fragments was selectively removed to increase specificity. Library preparation continued using the NEB Ultra II DNA Library Prep Kit (New England Biolabs, Ipswich, MA), with Illumina-compatible Y-adaptors and biotin-labeled DNA fragments were captured using streptavidin beads. To preserve the library’s complexity, the post-capture product was divided into two replicates, each with unique dual indices before PCR amplification. Sequencing was conducted at the Vincent J. Coates Genomics Sequencing Lab (Berkeley, CA) on an Illumina NovaSeq 6000 platform (Illumina, CA), generating approximately 2 million paired-end reads (2 x 150 bp) per gigabase (Gb) of genome size.

### HiFi SMRTbell library preparation

For long-read base sequence data, a HiFi SMRTbell library was constructed using the SMRTbell Express Template Prep Kit v2.0 (PacBio, Cat. #100-938-900) according to the manufacturer’s instructions. HMW gDNA was sheared to a target DNA size distribution between 15 and 18 kb. The sheared gDNA was concentrated using 0.45X of AMPure PB beads (PacBio, Cat. #100-265-900) for the removal of single-strand overhangs at 37°C for 15 minutes, followed by further enzymatic steps of DNA damage repair at 37°C for 30 minutes, end repair and A-tailing at 20°C for 10 minutes and 65°C for 30 minutes, and ligation of overhang adapter v3 at 20°C for 60 minutes. The SMRTbell library was purified and concentrated with 1X Ampure PB beads (PacBio, Cat. #100-265-900) for nuclease treatment at 37°C for 30 minutes and size selection using the BluePippin/PippinHT system (Sage Science, Beverly, MA; Cat #BLF7510/HPE7510) to collect fragments larger than 7-9 kb. The 15 – 20 kb average HiFi SMRTbell library was sequenced at UC Davis DNA Technologies Core (Davis, CA) using two 8M SMRT cells, Sequel II sequencing chemistry 2.0, and 30-hour movies, one on a PacBio Sequel II sequencer and one on a PacBio Sequel IIe sequencer.

### Nuclear Genome Assembly

To generate a high-contiguity assembly that preserves haplotypes, we assembled the genome of the bull kelp following the CCGP assembly pipeline for haploid species, as outlined in Table 1, which lists the tools and non-default parameters used in the assembly. The pipeline uses PacBio HiFi reads and Omni-C data to produce high-quality and highly contiguous genome assemblies. First, we removed the remnants adapter sequences from the PacBio HiFi dataset using HiFiAdapterFilt (Sim et al. 2022) and generated an initial haploid assembly using HiFiasm (Cheng et al. 2021, 2022), with the filtered PacBio HiFi reads and specifying no purging and the ploidy. From the generated output, we kept the file corresponding to the primary assembly file. We then aligned the Omni-C data to the assembly following the Arima Genomics Mapping Pipeline and then scaffolded it with SALSA (Ghurye et al. 2017, 2019).

**Table 1:** Sequencing and assembly statistics, and accession numbers.

|  |  |  |  |  |  |  |
| --- | --- | --- | --- | --- | --- | --- |
| Bio Projects & Vouchers | CCGP NCBI BioProject |  | PRJNA720569 |  |  |  |
|  | Genera NCBI BioProject |  | PRJNA765643 |  |  |  |
|  | Species NCBI BioProject |  | PRJNA777199 |  |  |  |
|  | NCBI BioSample |  | SAMN25872350 |  |  |  |
|  | Specimen identification |  | NB.12.F.B3 |  |  |  |
|  | NCBI Genome accessions |  |  |  |  |  |
|  | Assembly accession |  | JARUPZ000000000 |  |  |  |
|  | Genome sequence |  | GCA_031213475.1 |  |  |  |
| Genome Sequence | PacBio HiFi reads | Run | 1 PACBIO_SMRT (Sequel II) run: 1.5M spots, 21.9G bases, 13.6Gb, 1 PACBIO_SMRT (Sequel IIe) runs: 622,181 spots, 9.7G bases, 6.7Gb |  |  |  |
|  |  | Accession | SRX28564289, SRX28564290 |  |  |  |
|  | Omni-C Illumina reads | Run | 2 ILLUMINA (Illumina NovaSeq 6000) runs: 51.3M spots, 15.5G bases, 5.3G |  |  |  |
|  |  | Accession | SRX28564291-2 |  |  |  |
| Genome Assembly Quality Metrics | Assembly identifier (Quality code *) |  | uoNerLuet1(6.7.P6.Q57.C81) |  |  |  |
|  | HiFi Read coverage ** |  | 60.30X |  |  |  |
|  | Number of contigs |  | 2,121 |  |  |  |
|  | Contig N50 (bp) |  | 1,105,316 |  |  |  |
|  | Contig NG50 |  | 1,390,945 |  |  |  |
|  | Longest Contigs |  | 9,676,312 |  |  |  |
|  | Number of scaffolds |  | 1,562 |  |  |  |
|  | Scaffold N50 |  | 10,969,560 |  |  |  |
|  | Scaffold NG50 |  | 11,402,812 |  |  |  |
|  | Largest scaffold |  | 21,869,845 |  |  |  |
|  | Size of final assembly (bp) |  | 449,826,560 |  |  |  |
|  | Phased block NG50 § |  | 1,390,945 |  |  |  |
|  | Gaps per Gbp (#Gaps) |  | 1243(559) |  |  |  |
|  | Base pair QV |  | 57.822 |  |  |  |
|  | k-mer completeness |  | 84.9951 |  |  |  |
|  | BUSCO completeness** | C | S | D | F | M |
|  | (stramenopiles) n=100 | 72.00% | 69.00% | 3.00% | 12.00% | 16.00% |
| * Assembly quality code x.y.P.Q.C derived notation, from (Rhie et al. 2021). x = log10[contig NG50]; y = log10[scaffold NG50]; P = log10 [phased block NG50]; Q = Phred base accuracy QV (Quality value); C = % genome represented by the first ‘n’ scaffolds, following a known karyotype for this species of n=31(Genome on a Tree, query (*Nereocystis luetkeana*); Challis et al. 2023)<br>. Quality code for all the assembly denoted by primary assembly (uoNerLuet1.0.p) |  |  |  |  |  |  |
| § Read coverage and NGx statistics have been calculated based on the estimated genome size of 500 Mbp |  |  |  |  |  |  |
| ** BUSCO Scores. Complete BUSCOs (C). Complete and single-copy BUSCOs (S). Complete and duplicated BUSCOs (D). Fragmented BUSCOs (F). Missing BUSCOs (M). |  |  |  |  |  |  |

The assembly was manually curated by generating and analyzing their corresponding Omni-C contact maps and breaking scaffolds when misassemblies were identified. In general, to create the contact maps, we aligned the Omni-C data with BWA-MEM (Li 2013) and identified ligation junctions to generate Omni-C pairs (Lee et al. 2022) using pairtools (Open2C et al. 2023). We generated multi-resolution Omni-C matrices with cooler (Abdennur and Mirny 2020) and balanced them with hicExplorer (Ramírez et al. 2018). We used HiGlass (Kerpedjiev et al. 2018) and the PretextSuite to visualize the contact maps where we identified misassemblies and misjoins. Some remaining gaps were closed using the PacBio HiFi reads and YAGCloser. Finally, we checked for contamination using the BlobToolKit Framework (Challis et al. 2020) (Challis et al. 2020) and FCS-GX (Astashyn et al. 2023). In addition, we identified plastid scaffolds using HMMER 3.2 (Eddy 2011) by scanning for a set of HMMs of known marker genes.

### Genome quality assessment

We generated k-mer counts from the PacBio HiFi reads using *meryl*. The k-mer counts were then used in GenomeScope2.0 (Ranallo-Benavidez et al. 2020) to estimate genome features, including genome size, heterozygosity, and repeat content. To obtain general contiguity metrics, we ran QUAST (Gurevich et al. 2013). We used BUSCO (Manni et al. 2021) with Stramenopiles ortholog databases (stramenopiles_odb10) containing 100 genes to evaluate genome quality and functional completeness. Assessment of base level accuracy (quality value: QV) and k-mer completeness was performed using the previously generated *meryl* database and merqury (Rhie et al. 2020). Given the ploidy of the species, measurements of the size of the phased blocks are based on the size of the final contigs. We followed a quality metric nomenclature previously established (Rhie et al. 2021) with the genome quality code x.y.P.Q.C, where, x = log10[contig NG50]; y = log10[scaffold NG50]; P = log10 [phased block NG50]; Q = Phred base accuracy QV; C = % genome represented by the first ‘n’ scaffolds, following a karyotype of n=31 (Kemp 1960), estimated as the mode of the number of chromosomes from this species (Genome on a Tree, query= *Nereocystis luetkeana*; Challis et al. 2023).

### Mitochondrial Genome Assembly

We assembled the mitochondrial genome of *Nereocystis luetkeana* from the PacBio HiFi reads using the reference-guided pipeline MitoHiFi (Allio et al. 2020; Uliano-Silva et al. 2023), from which we also obtained the mitochondrial genome annotation. The mitochondrial sequence of an available *Nereocystis luetkeana* (NCBI:NC_042395.1; Zheng et al. 2019) was used as the starting sequence. After completion of the nuclear genome, we searched for matches of the resulting mitochondrial assembly sequence in the nuclear genome assembly using BLAST+ (Camacho et al. 2009) and filtered out contigs and scaffolds from the nuclear genome with a percentage of sequence identity > 99% and size smaller than the mitochondrial assembly sequence. No other manual curation was performed on the mitochondrial genome.

### Chloroplast Genome Assembly

The chloroplast assembly was done using the *Oatk* pipeline for *de novo* assembly of organelle genomes from PacBio HiFi data. We used the chloroplast genome assembly from *Saccharina japonica* (NCBI:NC_018523.1; Wang et al. 2013) as a guide for manual curation, in which we aligned the generated sequence against the guide, using *lastz* (Harris 2007), extracted contigs and fixed orientation when needed using samtools (Danecek et al. 2021) and *seqtk*. The alignment was visually validated using LAJ (Wilson et al. 2001). The resulting assembly was annotated using the online version of GeSeq (Tillich et al. 2017) and visualized using the online version of OGDRAW (Greiner et al. 2019).

### Annotation

The nuclear genome assembly of *Nereocystis luetkeana* was annotated using the JGI Annotation pipeline to produce a set of predicted gene models and their functional annotations (Grigoriev et al. 2014; Kuo et al. 2014). The pipeline is described briefly in the following steps. First, to account for repetitive sequences, the genome was masked using RepeatMasker (Smit et al. 1996) using the repeats from the RepBase library (Jurka et al. 2005) and the most frequently occurring repeats (>150 copies) identified by RepeatScout (Price et al. 2005). We applied multiple gene modelers to predict protein-coding genes, including *ab initio* modelers Fgenesh (Salamov and Solovyev 2000) and GeneMark (Ter-Hovhannisyan et al. 2008), homology-based Fgenesh+ and GeneWise (Birney et al. 2004) seeded by BLASTx alignments against the NCBI NR database, and transcriptome-based modelers Fgenesh and combest (Zhou et al. 2015). The transcriptome was assembled from the RNAseq reads using Trinity (v2.11.0)(Grabherr et al. 2011). Automated filtering based on homology and transcriptome support was applied to select the best representative gene model at each locus. Lastly, functional annotations of the predicted gene models were generated using various tools. SignalP 6.0 (Teufel et al. 2022) identified secretion signal sequences, TMHMM (Melén et al. 2003) detected transmembrane domains and InterProScan (Quevillon et al. 2005) identified protein domains. Homology searches were performed using Blastp against the NCBI NR, SwissProt, KEGG (Kanehisa et al. 2006), and TCDB (Saier et al. 2021) databases. The unique protein-coding genes are referenced in this manuscript by their protein identification number created by JGI. The genome assembly and annotations of *N. luetkeana* are also publicly available at the JGI PhycoCosm portal (Grigoriev et al. 2021).

### Orthogroup analysis

Translated non-redundant protein sequences of 10 phaeophyceae and 1 outgroup (*Schizocladia ischiensis*) reference genomes were obtained from the JGI PhycoCosm and Phaeoexplorer (Denoeud et al. 2024) portals, all taxa showing at least 70% BUSCO completeness provided by each database. The fasta files were included in OrthoFinder (Emms and Kelly 2015) to identify orthogroups with default parameters. A lineage-specific orthogroup was defined when only a single species, or group of species, possessed genes within that orthogroup. Orthogroup pairwise intersections and the enrichment of gene ontology terms in each orthogroup were made using R (R Core Team 2024) using packages ggplot2 (Wickham 2009), UpSetR (Conway et al. 2017) and topGO (Alexa and Rahnenfuhrer 2023).

## RESULTS

The Omni-C library generated 53.03 million read pairs and the PacBio HiFi library generated 2.15 million reads. The PacBio HiFi sequences yielded ∼80X genome coverage and had an N50 read length of 15,356 bp; a minimum read length of 230 bp; a mean read length of 14,681 bp; and a maximum read length of 51,198 bp (Supplementary Figure 1 for read length distribution). The coverage was estimated based on the estimates of genome size for other Laminariales (Kapraun 2005; Ye et al. 2015; Shan et al. 2020). The genome assembly size was calculated to be 449.77 mb. Based on the PacBio HiFi data, Genomescope 2.0 estimated a genome size of 416.29 Mb, and a 0.431% sequencing error rate. The k-mer spectrum shows an unimodal distribution with a major peak at ∼66-fold coverage (Figure 2A). The transcriptome showed 89% mapping to the genome, with 22,952 gene models predicted using JGI annotation pipeline with an average length of 8,312 bp and 5.89 exons per gene.

**Figure 1.**
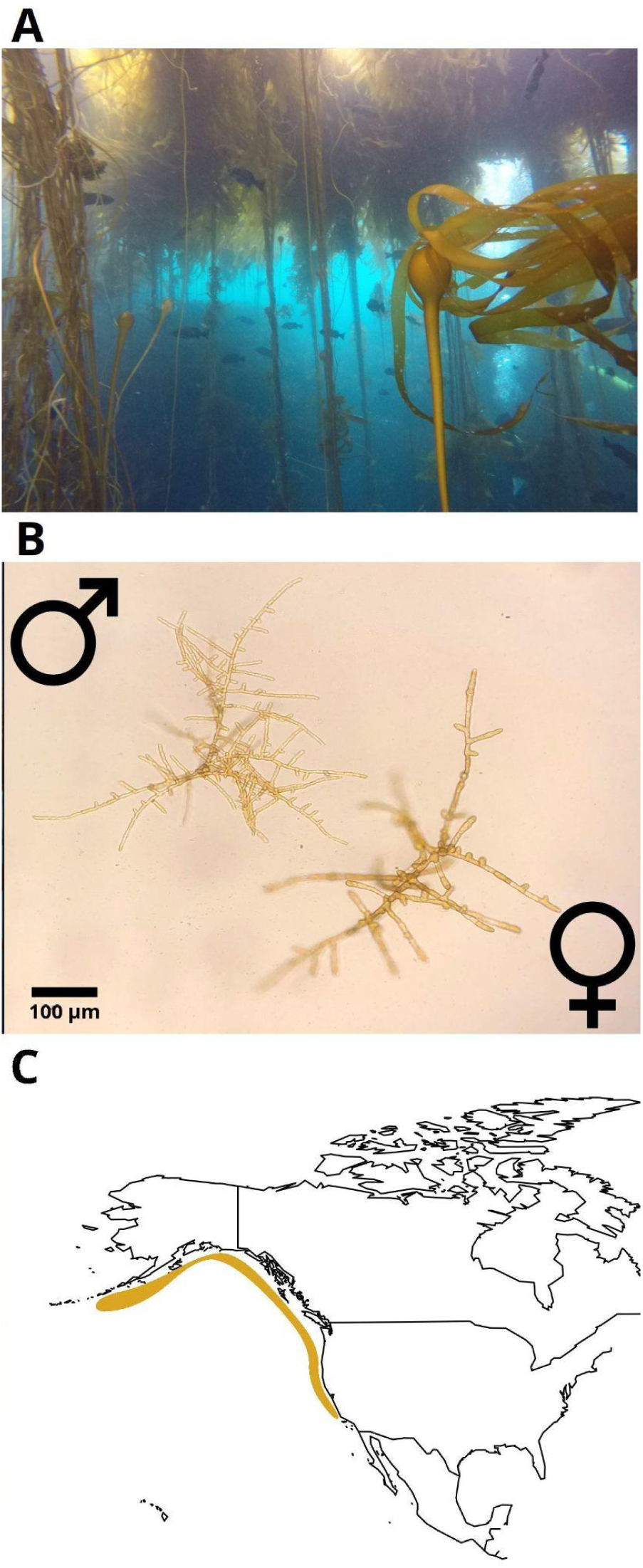
The bull kelp, *Nereocystis luetkeana*. A – Mature diploid sporophytes in the wild forming the marine forest canopy (Credit to Steve Lonhart). B) Female and male haploid microscopic gametophytes (Credit to Gabriel Montecinos). C – Distribution of *Nereocystis luetkeana* on Northeastern Pacific

**Figure 2.**
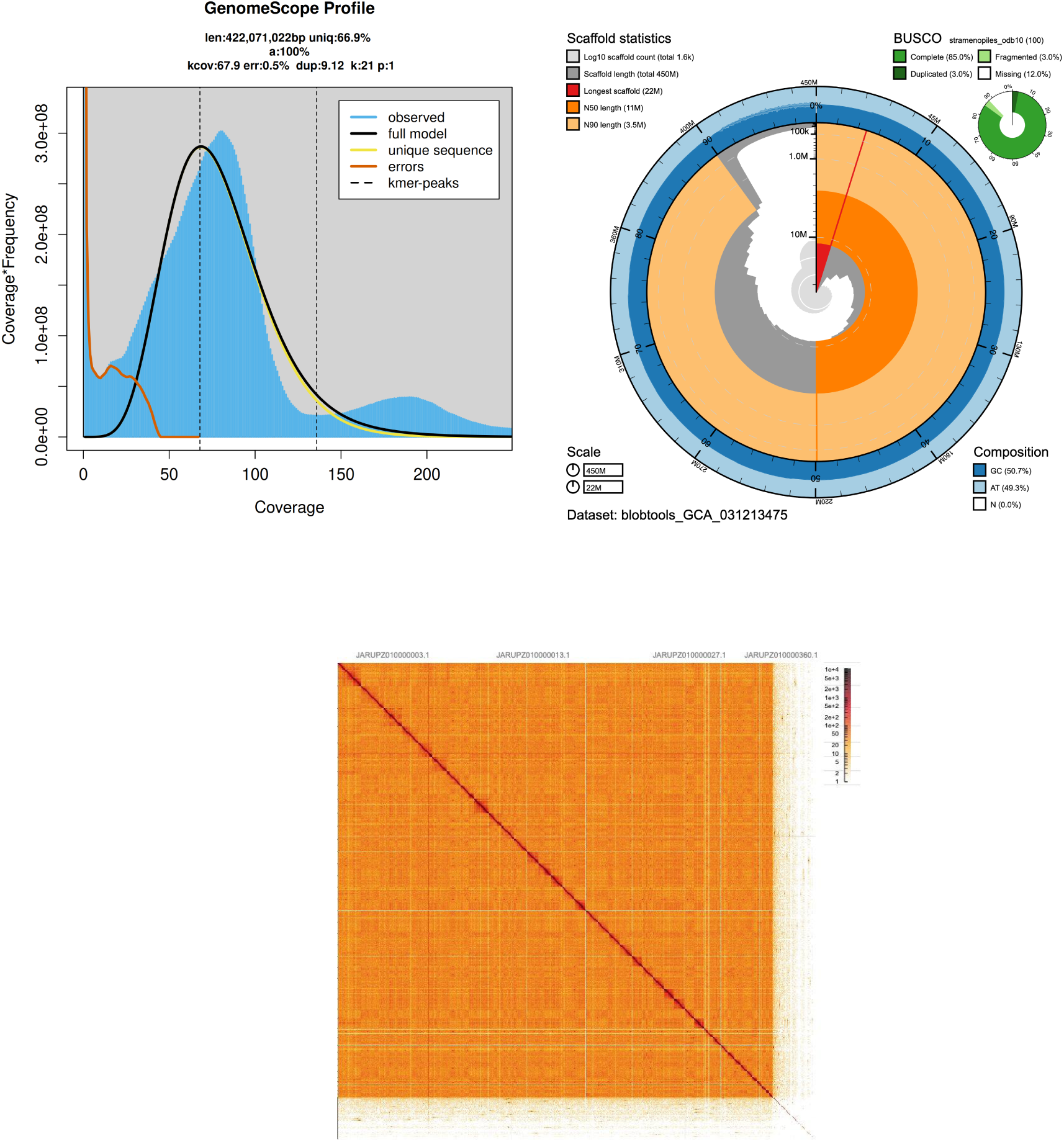
Visual overview of *Nereocystis luetkeana* genome assembly metrics and quality. A) A GenomeScope 2.0 k-mer spectrum from adapter-trimmed HiFi sequence data. B) BlobToolKit Snailplot showing N50 metrics for *N. luetkeana* assembly uoNerLuet1. Central circular plot shows genome length, with scaffolds added in clockwise fashion from longest to shortest. The outer ring indicates relative AT/CG content, and scaffold summary statistics and BUSCO scores for the stramenopiles_odb10 set of orthologues (n = 100). are shown at top left and right. C) Haploid contact map of uoNerLuet1 (10k resolution). Current data resolution: 640k.

### Nuclear genome assembly

The final genome assembly size (uoNerLuet1) is similar to the estimated genome assembly size from GenomeScope2.0. The assembly consists of 1,562 scaffolds spanning 449.82 Mb with a contig N50 of 1.1 Mb, a scaffold N50 of 10.96 Mb, the largest contig size of 9.67 Mb, and the largest scaffold size of 21.86 Mb. The assembly has a BUSCO completeness score for the Stramenopiles gene set of 72.0%, a base pair quality value (QV) of 57.82, and a kmer completeness of 84.99%.

During manual curation, we made 120 joins and 22 breaks based on the Omni-C contact map signal. We closed a single gap and we filtered out six contigs corresponding to mitochondrial contamination, 469 contigs corresponding to chloroplast contamination (cumulative size 17.83 Mb), and 525 (cumulative size 118.93Mb) corresponding to other sources of contamination (Supplementary Table 1). No other contigs were removed or modified. The Omni-C contact maps show a highly contiguous assembly with some chromosome-length scaffolds (Figure 2B). Assembly statistics are reported in Table 2 and represented graphically in Figure 2C. We have deposited the genome assembly on NCBI GenBank (see Table 2 and Data Availability for details).

**Table 2:** Assembly Pipeline and Software Used. Software citations are listed in the text.

| Assembly | Software and any non-default options | Version | Reference |
| --- | --- | --- | --- |
| Filtering PacBio HiFi adapters | HiFiAdapterFilt | Commit 64d1c7b | Sim et al. 2022 |
| K-mer counting | Meryl (k=31) | 1 | https://github.com/marbl/meryl |
| Estimation of genome size and heterozygosity | GenomeScope (k=31) | 2 | Ranallo-Benavidez et al. 2020 |
| *De novo assembly (contiging)* | HiFiasm (-primary, -l0, output p_ctg) | 0.16.1-r375 | Cheng et al. 2021<br>Cheng et al. 2022 |
| **Scaffolding** |  |  |  |
| Omni-C data alignment | <u>Arima Genomics Mapping Pipeline</u> | Commit 2e74ea4 | https://github.com/ArimaGenomics/mapping_pipeline |
| <u>Arima Genomics Mapping Pipeline (AGMP)</u> | BWA-MEM | 0.7.17-r1188 | Li 2013 |
|  | samtools | 1.11 | Danecek et al. 2021 |
|  | filter_five_end.pl (AGMP) | Commit 2e74ea4 | https://github.com/ArimaGenomics/mapping_pipeline |
|  | two_read_bam_combiner.pl (AGMP) | Commit 2e74ea4 | https://github.com/ArimaGenomics/mapping_pipeline |
|  | picard | 2.27.5 | https://broadinstitute.github.io/picard/ |
| Omni-C Scaffolding | SALSA (-DNASE, -i 20, -p yes) | 2 | Ghurye et al. 2017, Ghurye et al. 2019 |
| Gap closing | YAGCloser (-mins 2 -f 20 -mcc 2 -prt 0.25 -eft 0.2 -pld 0.2) | Commit 0e34c3b | http://github.com/merlyescalona/yagcloser |
| **Omni-C Contact map generation** |  |  |  |
| Short-read alignment | BWA-MEM (-5SP) | 0.7.17-r1188 | Li 2013 |
| SAM/BAM processing | samtools | 1.11 | Danecek et al. 2021 |
| SAM/BAM filtering | pairtools | 0.3.0 | Open2C et al. 2024 |
| Pairs indexing | pairix | 0.3.7 | Lee et al. 2022 |
| Matrix generation | cooler | 0.8.10 | Abdennur and Mirny 2020 |
| Matrix balancing | hicExplorer (hicCorrectmatrix correct --filterThreshold -2 4) | 3.6 | Ramírez et al. 2018 |
| Contact map visualization | HiGlass | 2.1.11 | Kerpedjiev et al. 2018 |
|  | PretextMap | 0.1.4 | https://github.com/wtsi-hpag/PretextView |
|  | PretextView | 0.1.5 | https://github.com/wtsi-hpag/PretextMap |
|  | PretextSnapshot | 0.0.3 | https://github.com/wtsi-hpag/PretextSnapshot |
| Manual curation tools | Rapid curation pipeline (Wellcome Trust Sanger Institute, Genome Reference Informatics Team) | Commit 7acf220c | https://gitlab.com/wtsi-grit/rapid-curation |
| **Genome quality assessment** |  |  |  |
| Basic assembly metrics | QUAST (--est-ref-size) | 5.0.2 | Gurevich et al. 2013 |
| Assembly completeness | BUSCO (-m geno, -l stramenopiles) | 5.0.0 | Manni et al. 2021 |
|  | Mercury | 2020-01-29 | Rhie et al. 2020 |
| **Contamination screening** |  |  |  |
| Local alignment tool | BLAST+ (-db nt, -outfmt '6 qseqid staxids bitscore std' , -max_target_seqs 1, -max_hsps 1, -evalue 1e-25 ) | 2.15 | Camacho et al. 2009 |
| General contamination screening | BlobToolKit (PacBio HiFi Coverage, NCBI Taxa ID = 117523, BUSCODB = stramenopiles) | 2.3.3 | Challis et al. 2020 |
| **Mitochondrial assembly** |  |  |  |
| Mitochondrial genome assembly | MitoHiFi (-r, -p 80, -o 1, -a plant) Reference: Nereocystis luetkeana (NCBI:NC_042395.1) | 3 | Uliano-Silva et al. 2023 |
| Genome annotation | MitoFinder (MitoHiFi pipeline parameters) | 1.4 | Allio et al. 2020 |
| **Chloroplast assembly** |  |  |  |
| de novo genome assembly | Oatk (syncasm -k 1001 -c 400) | 1 | https://github.com/c-zhou/oatk |
| Sequence editing | seqtk | 1.3-r115-dirty | https://github.com/lh3/seqtk |
| Sequence alignment | lastz (--nogapped,--notransition, | 1.04.15 | Harris 2017 |
|  | –step=20) |  |  |
| Alignment<br>visualization | LAJ<br>(http://globin.cse.psu.edu/dist/laj/) | 2005-12-14 | Wilson et al. 2001 |
| Genome annotation | GeSeq<br>(https://chlorobox.mpimp-golm.mpg.de/geseq.html) | 2021 | Tillich et al. 2017 |

### Mitochondrial genome assembly

We assembled a partial mitochondrial genome for *Nereocystis luetkeana* with MitoHiFi. The final mitochondrial sequence is 37,397 bp long, with base composition of A=35.92%, C=9.418%, G=14.99%, and T=39.67%. It consists of 2 rRNAs, 23 unique transfer RNAs, and 38 protein-coding genes.

### Chloroplast genome assembly

The final chloroplast assembly of *N. luetkeana* is 131,016 bp long and has 31.12% GC content. The large single copy (LSC) is 78,062 bp long, the small single copy (SSC) is 45,058 bp long, and the inverted repeat is 3,933 bp long. This plastome contains 85 genes, including rRNAs and tRNAs (not counting the duplication of the IR).

### Orthogroup analysis and Gene Ontology enrichment

The OrthoFinder analysis revealed 4,762 orthogroups common to all 11 taxa (Figure 5), of which 850 are common to all analyzed phaeophyceae. *N. luetkeana* and *M. pyrifera* had more lineage-specific orthogroups (542) than any other species or group intersection. These orthogroups comprised 858 and 907 genes, of which 320 and 200 had known annotations in *N. luetkeana* and *M. pyrifera,* respectively (Table 2). From these, the molecular function “ATP binding” was the most enriched GO term in *M. pyrifera,* with 21 ATPase genes and 10 in *N. luetkeana*. These enriched ATPases were classified as dynein-related (IPR011704), Na^+^-K^+^ transport P-type ATPase (IPR004014), and AAA-type (IPR003960) in both kelps. No cellular components nor biological processes were significantly enriched.

**Figure 3.**
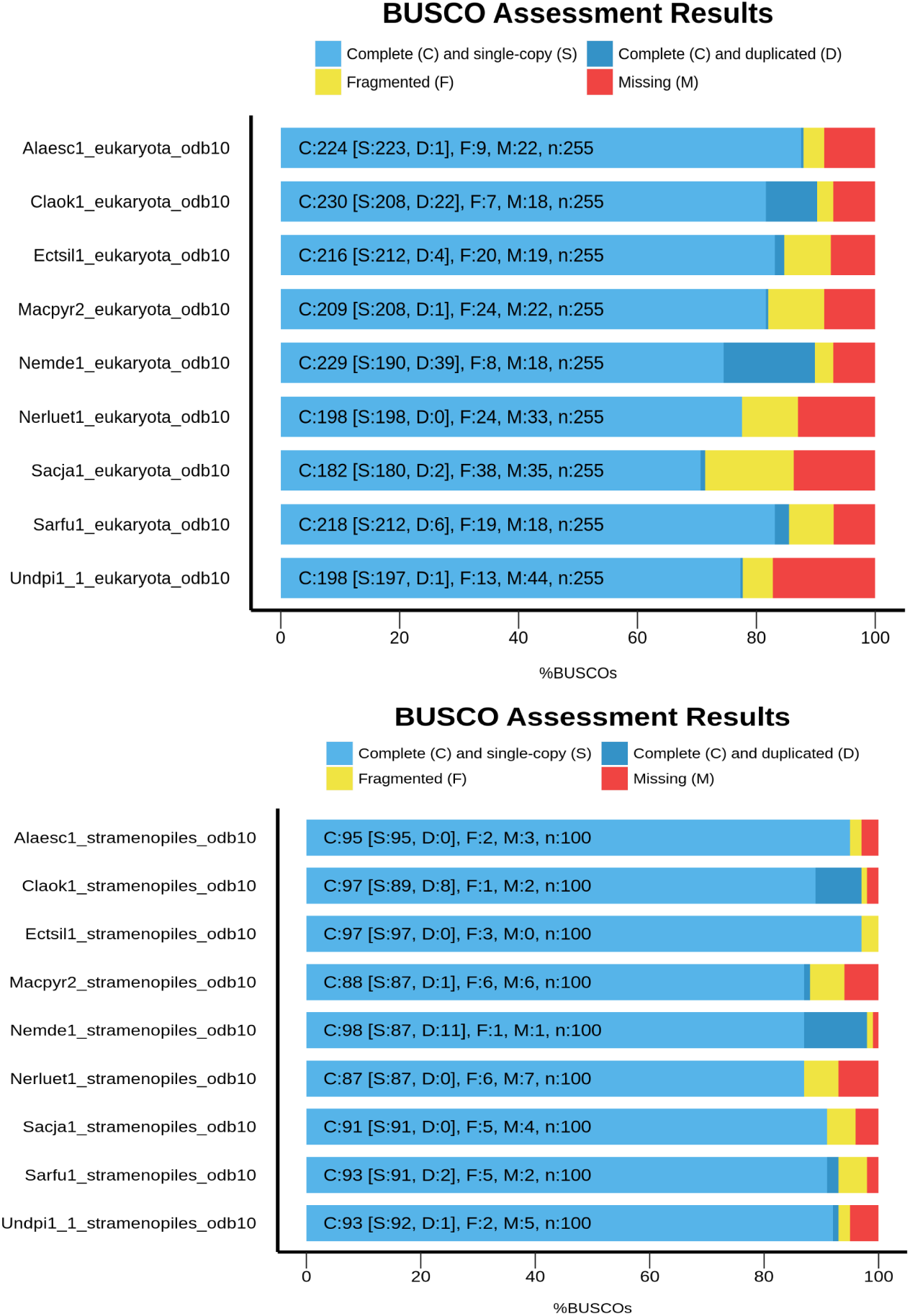
BUSCO’S eukaryote (n=255) and stramenopiles (n = 100) ortholog gene groups set on *Nereocystis luetkeana (Nerluet1)* assembly and other phaeophyceae seaweed genome assemblies, specifically: *Alaesc: Alaria esculenta, Cloak: Cladosiphon okamuranus, Ectsil: Ectocarpus siliculosus, Macpyr2: Macrocystis piryfera, Nemde1: Nemacystus decipiens, Sacja: Saccharina japonica, Sarfu: Sargassum fusiforme, Undpi1: Undaria pinnatifica*.

**Figure 4.**
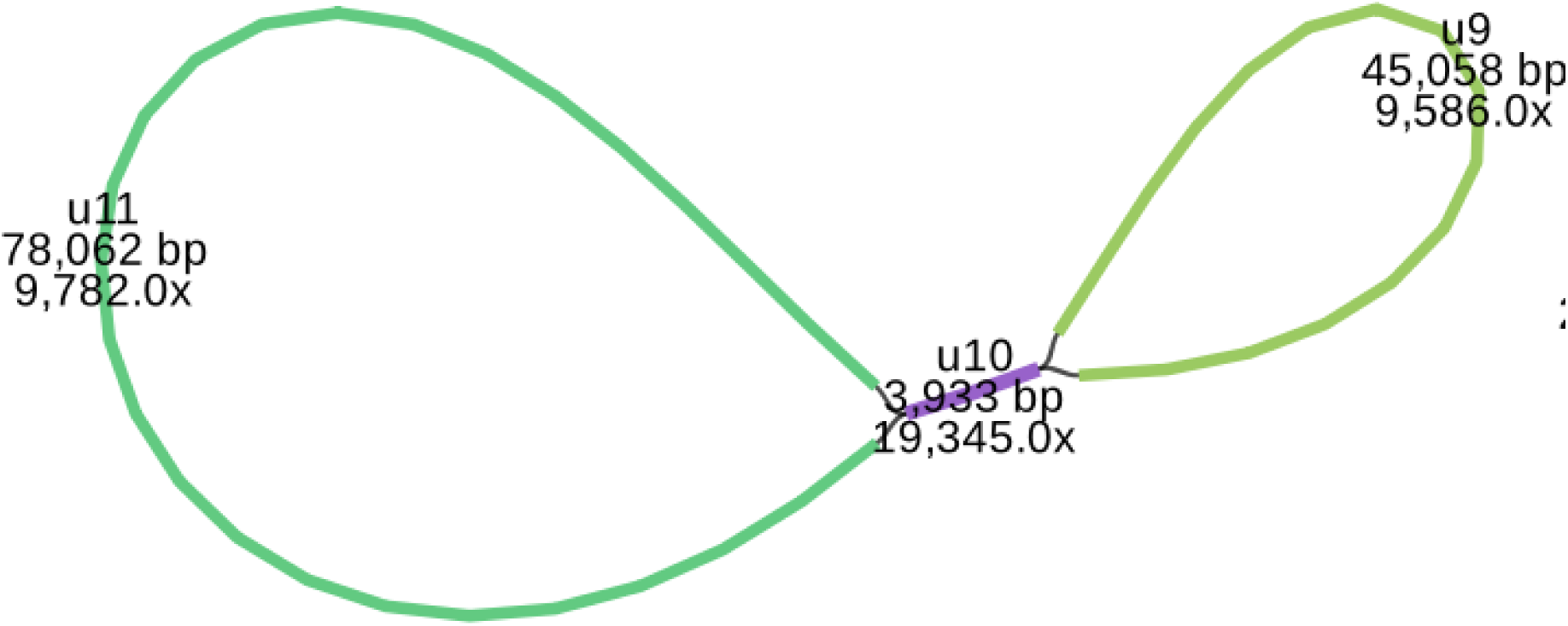
Model of the putative chloroplast structure for *Nereocystis luetkeana*.

**Figure 5.**
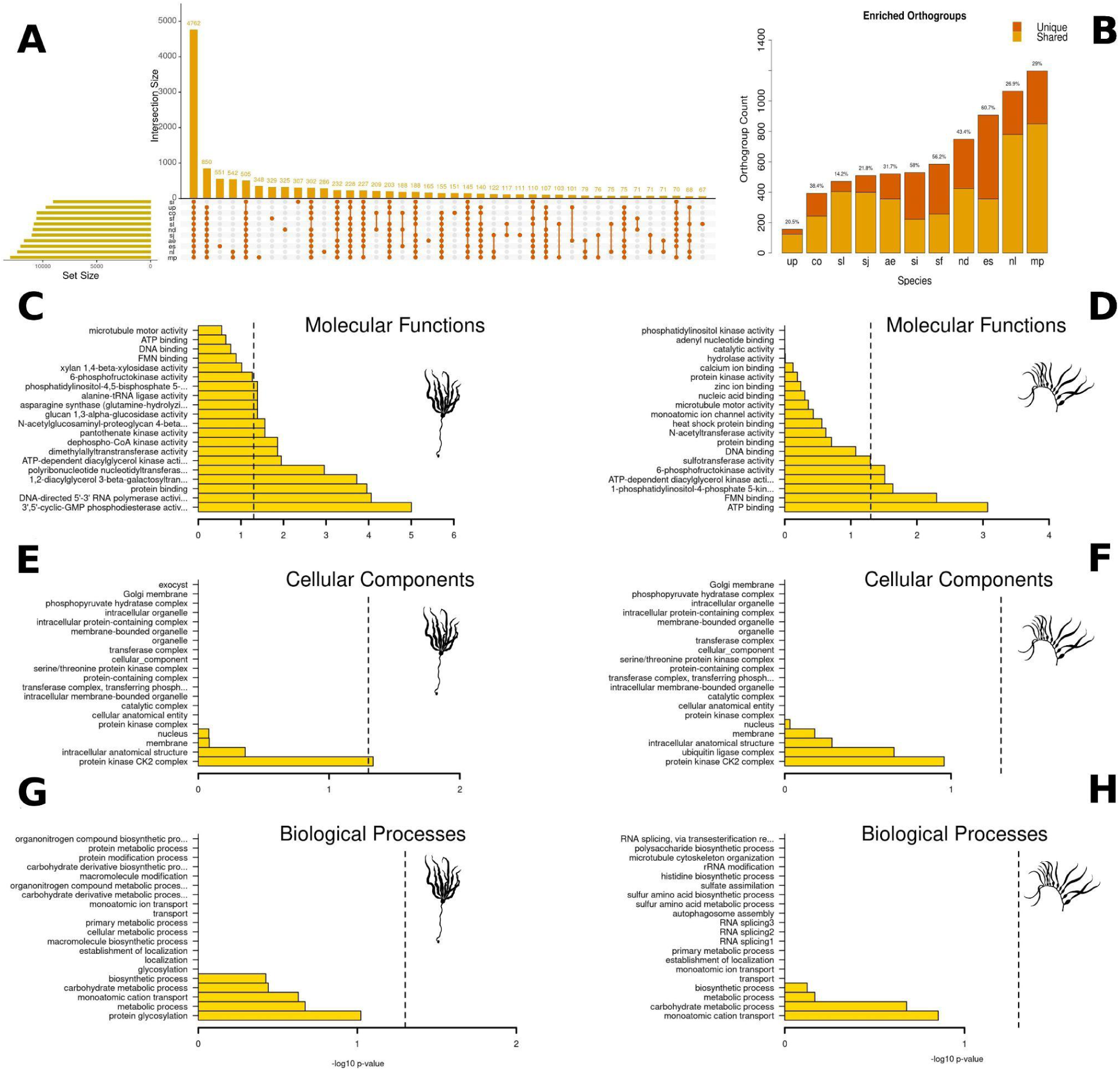
Comparative orthogroup and gene ontology enrichment analysis of *Nereocystis luetkeana* genome. In A, an upset plot showing all orthogroup counts for intersection groups between 10 other phaeophyceae genomes. B) The expanded orthogroups in each species, highlighting the proportion of unique orthogroups. In C, D, E, F, G and H, enrichment analysis of gene ontology terms of the 592 unique orthogroup genes present in both *N. luetkeana* (left) and *M. pyrifera* (right). C and D) Molecular functions; E and F) Cellular Components; G and H) Biological Processes. The acronyms are: “ae”: *Alaria esculenta*,”co”: *Cladosiphon okamuranus*, “es”: *Ectocarpus siliculosus*, “mp”: *Macrocystis pyrifera*, “nd”: *Nemacystus decipiens*, “sf”: *Sargassum fusiforme,* “si”: *Schizocladia ischiensis* (outgroup), “sj”: *Saccharina japonica*,“sl”: *Saccharina latissima*, “up”: *Undaria pinnatifida*. The silhouette images were obtained at https://www.phylopic.org/, credit of *Macrocystis* silhouette to Harold N Eyster, 2019.

## DISCUSSION

Considering seaweed diversity, available reference genomes are underrepresented compared to land plants. Almost half of vascular plant orders have at least one complete genome available (Marks et al. 2021), while only 32% (n=16) of multicellular algae orders are represented (Cock et al. 2010; Collén et al. 2013; Brawley et al. 2017; Gabrielson et al. 2018; De Clerck et al. 2018; Kinnby et al. 2020; Thapa et al. 2020; Graf et al. 2021; Steele et al. 2022; Nakamura-Gouvea et al. 2022). Recently, 60 new genomes of Phaeophyceae have been sequenced in a considerable effort by the Phaeoexplorer project (Denoeud et al. 2024), which provided 14 new families sequenced and 17 good-quality genomes. The giant kelp*, Macrocystis pyrifera,* has also been sequenced to the chromosome level (Diesel et al. 2023). The CCGP initiative is contributing to expanding the available seaweed genomes by sequencing the monospecific kelp genus *Nereocystis*.

We provide a high-quality genome of *N. luetkeana* compared to the quality of currently available kelp genomes. *N. luetkeana* genome is larger than the average Phaeophyceae genomes (321 mb) but slightly smaller (10%) than other Laminariales (496 mb), as previously reported from DAPI fluorimetry (Kapraun 2005). The number of detected genes is similar to *M. pyrifera* but 30% larger than *Sacharina japonica.* The genome of *Laminaria digitata* could not be compared because it is highly fragmented (Denoeud et al. 2024). As important marine forest-forming species that sustain significant biodiversity, more genomes from members of the Laminariaceae family are needed as genomics is becoming an essential conservation biology tool (Fiedler et al. 2022; Theissinger et al. 2023).

The enrichment of specific orthogroups for *N. luetkeana* and *M. pyrifera* can be related to their large body size and the new gene functions needed to sustain their remarkable development. The presence of unique ATP-binding protein-coding genes in both *N. luetkeana* and *M. pyrifera* may be linked to the increased metabolic flux required by their rapid growth rate. Similarly, the elevated energetic demands for synthesizing carbon-rich molecules have already been described in other large body-sized organisms. In birds and mammals, Na^+^-K^+^-ATPases gene enrichment and higher enzymatic activity have been associated with body size changes of three orders of magnitude, primarily associated with lipid biosynthesis (Turner et al. 2005). For photosynthetic eukaryotes, ATPase upregulation has been associated with higher growth and rubisco protein levels in *Ulva lactuca* (El-Adl et al. 2021) and increased nitrogen assimilation and overall yield in *Oryza sativa* (Zhang et al. 2021). The synthesis of polysaccharides, such as alginate and fucoidan, and their transport via cytoskeletal trafficking of Golgi-derived vesicles require elevated ATP hydrolysis (Nagasato and Motomura 2009; Cosgrove 2024). This is crucial for cell wall thickening during the rapid cell division and elongation in basal meristems in the massive sporophytes (Bisgrove and Kropf 2001; Starko et al. 2018). Furthermore, the enrichment of ATPases in large-body kelps is consistent with earlier findings of high ATP concentration in sieve elements (Schmitz and Srivastava 1974). This supports the energy-intensive process of rapid, long-distance translocation of photoassimilates, essential for sustaining the growth of these large kelps (Schmitz and Srivastava 1974; Schmitz and Lobban 1976; Knoblauch et al. 2016a).

The association between ATP-binding protein-coding genes and body size could be further detailed when the closest relatives to bull kelp, *Postelsia palmaeformis*, genome is sequenced given its much smaller body size. Conversely, another large-body, closely related species is the elk kelp, *Pelagophycus porra* (Parker and Bleck 1965), an understudied, ironically cryptic species (deep habitat) with no genomic resources. The genomes of two other large-bodied kelps from related groups, the 25-meter-long kelp *Eualaria fistulosa* (Alariaceae) (Setchell and Gardner 1925) and the 17.5-meter-long *Himantothallus grandifolius* (Desmarestiaceae) (Dieckmann et al. 1985), could also provide important insights into gene repertoire evolution responsible for kelp body size.

The genomic resources developed here enable a suite of *narrow-sense* population genomics tools to assist the species’ conservation biology (Luikart et al. 2018). For example, high-density SNP panels mapped to this genome will allow pinpointing specific genomic features when detecting signatures of selection and their association with environmental variation. Likewise, transcriptomics experiments (RNAseq) can now produce sets of differentially expressed genes and test their enrichment in different conditions. Patterns of linkage disequilibrium decay across the genome can now be characterized and used in demographic inference.

## Acknowledgements

PacBio Sequel II/IIe library prep and sequencing was carried out at the DNA Technologies and Expression Analysis Core at the UC Davis Genome Center, supported by NIH Shared Instrumentation Grant 1S10OD010786-01. Deep sequencing of Omni-C libraries used the Novaseq S4 sequencing platforms at the Vincent J. Coates Genomics Sequencing Laboratory at UC Berkeley, supported by NIH S10 OD018174 Instrumentation Grant. We thank the staff at the UC Davis DNA Technologies and Expression Analysis Core and the UC Santa Cruz Paleogenomics Laboratory for their diligence and dedication to generating high-quality sequence data. We thank Dr. Courtney Miller, and Dr. Erin Toffelmier for help with coordination, sample submission and interacting with the CCGP community.

## Data Availability

Data generated for this study are available under NCBI BioProject PRJNA777199. Raw sequencing data for samples NB.12.F.B3 (NCBI BioSamples SAMN25872350) are deposited in the NCBI Short Read Archive (SRA) under PENDING for PacBio HiFi sequencing data, and PENDING for the Omni-C Illumina sequencing data. GenBank accession for the assembly is GCA_031213475.1; and for annotated genome JARUPZ000000000. Assembly scripts and other data for the analyses presented can be found at the following GitHub repository: www.github.com/ccgproject/ccgp_assembly

## REFERENCES

1. Abdennur N, Mirny LA (2020) Cooler: scalable storage for Hi-C data and other genomically labeled arrays. Bioinforma Oxf Engl 36:311–316. 10.1093/bioinformatics/btz540

2. Ahn O, Petrell R, Harrison P (1998) Ammonium and nitrate uptake by *Laminaria saccharina* and *Nereocystis luetkeana* originating from a salmon sea cage farm. J Appl Phycol 10:333–340. 10.1023/A:1008092521651

3. Alexa A, Rahnenfuhrer J (2023) topGO: Enrichment Analysis for Gene Ontology.

4. Allio R, Schomaker-Bastos A, Romiguier J, et al (2020) MitoFinder: Efficient automated large-scale extraction of mitogenomic data in target enrichment phylogenomics. Mol Ecol Resour 20:892–905. 10.1111/1755-0998.13160

5. Amsler CD, Neushul M (1989) Diel periodicity of spore release from the kelp *Nereocystis luetkeana* (Mertens) Postels et Ruprecht. J Exp Mar Biol Ecol 134:117–127. 10.1016/0022-0981(90)90104-K

6. Assis J, Araújo MB, Serrão EA (2018) Projected climate changes threaten ancient refugia of kelp forests in the North Atlantic. Glob Change Biol 24:e55–e66. 10.1111/gcb.13818

7. Astashyn A, Tvedte ES, Sweeney D, et al (2023) Rapid and sensitive detection of genome contamination at scale with FCS-GX. BioRxiv Prepr Serv Biol 2023.06.02.543519. 10.1101/2023.06.02.543519

8. Bemmels JB, Starko S, Weigel BL, et al (2025) Population genomics reveals strong impacts of genetic drift without purging and guides conservation of bull and giant kelp. Curr Biol 35:688–698.e8. 10.1016/j.cub.2024.12.025

9. Berry HD, Mumford TF, Christiaen B, et al (2021) Long-term changes in kelp forests in an inner basin of the Salish Sea. PLOS ONE 16:e0229703. 10.1371/journal.pone.0229703

10. Birney E, Clamp M, Durbin R (2004) GeneWise and Genomewise. Genome Res 14:988–995. 10.1101/gr.1865504

11. Bisgrove SR, Kropf DL (2001) Cell wall deposition during morphogenesis in fucoid algae. Planta 212:648–658. 10.1007/s004250000434

12. Brawley SH, Blouin NA, Ficko-Blean E, et al (2017) Insights into the red algae and eukaryotic evolution from the genome of *Porphyra umbilicalis* (Bangiophyceae, Rhodophyta). Proc Natl Acad Sci U S A 114:E6361–E6370. 10.1073/pnas.1703088114

13. Breen PA, Miller DC, Adkins BE (1976) An Examination of Harvested Sea Urchin Populations in the Tofino Area. Fisheries Research Board of Canada, Pacific Biological Station, Nanaimo, B.C.

14. Burnett NP, Ricart AM, Winquist T, et al (2024) Bimodal spore release heights in the water column enhance local retention and population connectivity of bull kelp, Nereocystis luetkeana. Ecol Evol 14:e70177. 10.1002/ece3.70177

15. Camacho C, Coulouris G, Avagyan V, et al (2009) BLAST+: architecture and applications. BMC Bioinformatics 10:421. 10.1186/1471-2105-10-421

16. Challis R, Kumar S, Sotero-Caio C, et al (2023) Genomes on a Tree (GoaT): A versatile, scalable search engine for genomic and sequencing project metadata across the eukaryotic tree of life. Wellcome Open Res 8:24. 10.12688/wellcomeopenres.18658.1

17. Challis R, Richards E, Rajan J, et al (2020) BlobToolKit – Interactive Quality Assessment of Genome Assemblies. G3 GenesGenomesGenetics 10:1361–1374. 10.1534/g3.119.400908

18. Cheng H, Concepcion GT, Feng X, et al (2021) Haplotype-resolved de novo assembly using phased assembly graphs with hifiasm. Nat Methods 18:170–175. 10.1038/s41592-020-01056-5

19. Cheng H, Jarvis ED, Fedrigo O, et al (2022) Haplotype-resolved assembly of diploid genomes without parental data. Nat Biotechnol 40:1332–1335. 10.1038/s41587-022-01261-x

20. Cock JM, Sterck L, Rouzé P, et al (2010) The Ectocarpus genome and the independent evolution of multicellularity in brown algae. Nature 465:617–621. 10.1038/nature09016

21. Collén J, Porcel B, Carré W, et al (2013) Genome structure and metabolic features in the red seaweed *Chondrus crispus* shed light on evolution of the Archaeplastida. Proc Natl Acad Sci 110:5247–5252. 10.1073/pnas.1221259110

22. Conway JR, Lex A, Gehlenborg N (2017) UpSetR: an R package for the visualization of intersecting sets and their properties. Bioinformatics 33:2938–2940. 10.1093/bioinformatics/btx364

23. Cosgrove DJ (2024) Structure and growth of plant cell walls. Nat Rev Mol Cell Biol 25:340–358. 10.1038/s41580-023-00691-y

24. Danecek P, Bonfield JK, Liddle J, et al (2021) Twelve years of SAMtools and BCFtools. GigaScience 10:giab008. 10.1093/gigascience/giab008

25. Dayton PK (1985) Ecology of Kelp Communities. Annu Rev Ecol Syst 16:215–245

26. Dayton PK, Currie V, Gerrodette T, et al (1984) Patch Dynamics and Stability of Some California Kelp Communities. Ecol Monogr 54:253–289. 10.2307/1942498

27. De Clerck O, Kao S-M, Bogaert KA, et al (2018) Insights into the evolution of multicellularity from the sea lettuce genome. Curr Biol 28:2921–2933.e5. 10.1016/j.cub.2018.08.015

28. Denny M, Gaylord B, Cowen E (1997) Flow and flexibility – II. The roles of size and shape in determining wave forces on the bull kelp *Nereocystis luetkeana*. J Exp Biol 200:3165–3183

29. Denoeud F, Godfroy O, Cruaud C, et al (2024) Evolutionary genomics of the emergence of brown algae as key components of coastal ecosystems. 2024.02.19.579948

30. Dieckmann G, Reichardt W, Zieliński K (1985) Growth and Production of the Seaweed, Himantothallus grandifolius, at King George Island. In: Siegfried WR, Condy PR, Laws RM (eds) Antarctic Nutrient Cycles and Food Webs. Springer, Berlin, Heidelberg, pp 104–108

31. Diesel J, Molano G, Montecinos GJ, et al (2023) A scaffolded and annotated reference genome of giant kelp (Macrocystis pyrifera). BMC Genomics 24:543. 10.1186/s12864-023-09658-x

32. Druehl LD (1970) The pattern of Laminariales distribution in the northeast Pacific. Phycologia 9:237–247. 10.2216/i0031-8884-9-3-237.1

33. Duncan MJ, Foreman RE (1980) Phytochrome-mediated stipe elongation in the kelp *Nereocystis luetkeana*. J Phycol 16:138–142. 10.1111/j.1529-8817.1980.tb03008.x

34. Eddy SR (2011) Accelerated Profile HMM Searches. PLoS Comput Biol 7:e1002195. 10.1371/journal.pcbi.1002195

35. El-Adl MF, El-Katony TM, Nada RM (2021) High external Na+, but not K+, stimulates the growth of *Ulva lactuca* (L.) via induction of the plasma membrane *ATPases* and achievement of K+/Na+ homeostasis. Plant Physiol Biochem 163:239–249. 10.1016/j.plaphy.2021.03.032

36. Emms DM, Kelly S (2015) OrthoFinder: solving fundamental biases in whole genome comparisons dramatically improves orthogroup inference accuracy. Genome Biol 16:157. 10.1186/s13059-015-0721-2

37. Fiedler PL, Erickson B, Esgro M, et al (2022) Seizing the moment: The opportunity and relevance of the California Conservation Genomics Project to state and federal conservation policy. J Hered 113:589–596. 10.1093/jhered/esac046

38. Finger DJI, McPherson ML, Houskeeper HF, Kudela RM (2021) Mapping bull kelp canopy in northern California using Landsat to enable long-term monitoring. Remote Sens Environ 254:112243. 10.1016/j.rse.2020.112243

39. Foreman RE (1977) Benthic community modification and recovery following intensive grazing by *Strongylocentrotus droebachiensis*. Helgoländer Wiss Meeresunters 30:468–484. 10.1007/BF02207855

40. Foreman RE (1970) Physiology, ecology, and development of the brown alga, Nereocystis luetkeana (Mertens) P. & R. Ph.D., University of California, Berkeley

41. Gabrielson PW, Hughey JR, Diaz-Pulido G (2018) Genomics reveals abundant speciation in the coral reef building alga *Porolithon onkodes* (Corallinales, Rhodophyta). J Phycol 54:429–434. 10.1111/jpy.12761

42. Ghurye J, Pop M, Koren S, et al (2017) Scaffolding of long read assemblies using long range contact information. BMC Genomics 18:527. 10.1186/s12864-017-3879-z

43. Ghurye J, Rhie A, Walenz BP, et al (2019) Integrating Hi-C links with assembly graphs for chromosome-scale assembly. PLoS Comput Biol 15:e1007273. 10.1371/journal.pcbi.1007273

44. Gierke L, Coelho NC, Khangaonkar T, et al (2023) Range wide genetic differentiation in the bull kelp Nereocystis luetkeana with a seascape genetic focus on the Salish Sea. Front Mar Sci 10:. 10.3389/fmars.2023.1275905

45. Grabherr MG, Haas BJ, Yassour M, et al (2011) Full-length transcriptome assembly from RNA-Seq data without a reference genome. Nat Biotechnol 29:644–652. 10.1038/nbt.1883

46. Graf L, Shin Y, Yang JH, et al (2021) A genome-wide investigation of the effect of farming and human-mediated introduction on the ubiquitous seaweed *Undaria pinnatifida*. Nat Ecol Evol 5:360–368. 10.1038/s41559-020-01378-9

47. Greiner S, Lehwark P, Bock R (2019) OrganellarGenomeDRAW (OGDRAW) version 1.3.1: expanded toolkit for the graphical visualization of organellar genomes. Nucleic Acids Res 47:W59–W64. 10.1093/nar/gkz238

48. Grigoriev IV, Hayes RD, Calhoun S, et al (2021) PhycoCosm, a comparative algal genomics resource. Nucleic Acids Res 49:D1004–D1011. 10.1093/nar/gkaa898

49. Grigoriev IV, Nikitin R, Haridas S, et al (2014) MycoCosm portal: gearing up for 1000 fungal genomes. Nucleic Acids Res 42:D699–704. 10.1093/nar/gkt1183

50. Gurevich A, Saveliev V, Vyahhi N, Tesler G (2013) QUAST: quality assessment tool for genome assemblies. Bioinforma Oxf Engl 29:1072–1075. 10.1093/bioinformatics/btt086

51. Hamilton SL, Bell TW, Watson JR, et al (2020) Remote sensing: generation of long-term kelp bed data sets for evaluation of impacts of climatic variation. Ecology 101:1–13

52. Harris RS (2007) Improved pairwise alignment of genomic dna. Phd, Pennsylvania State University

53. Inglis PW, Pappas M de CR, Resende LV, Grattapaglia D (2018) Fast and inexpensive protocols for consistent extraction of high quality DNA and RNA from challenging plant and fungal samples for high-throughput SNP genotyping and sequencing applications. PloS One 13:e0206085. 10.1371/journal.pone.0206085

54. Jurka J, Kapitonov VV, Pavlicek A, et al (2005) Repbase Update, a database of eukaryotic repetitive elements. Cytogenet Genome Res 110:462–467. 10.1159/000084979

55. Kain J (1987) Patterns of relative growth in *Nereocystis luetkeana* (Phaeophyta). J Phycol 23:181–187. 10.1111/j.0022-3646.1987.00181.x

56. Kain J, Norton T (1987) Growth of blades of *Nereocystis luetkeana* (Phaeophyta) in darkness. J Phycol 23:464–469. 10.1111/j.1529-8817.1987.tb02533.x

57. Kanehisa M, Goto S, Hattori M, et al (2006) From genomics to chemical genomics: new developments in KEGG. Nucleic Acids Res 34:D354–357. 10.1093/nar/gkj102

58. Kapraun DF (2005) Nuclear DNA content estimates in multicellular green, red and brown algae: phylogenetic considerations. Ann Bot 95:7–44. 10.1093/aob/mci002

59. Kemp CL (1960) Chromosomal alternation of generations in Nereocystis luetkeana (mertens) Postels and Ruprecht. University of British Columbia

60. Kerpedjiev P, Abdennur N, Lekschas F, et al (2018) HiGlass: web-based visual exploration and analysis of genome interaction maps. Genome Biol 19:125. 10.1186/s13059-018-1486-1

61. Kinnby A, Jonsson PR, Ortega-Martinez O, et al (2020) Combining an Ecological Experiment and a Genome Scan Show Idiosyncratic Responses to Salinity Stress in Local Populations of a Seaweed. Front Mar Sci 7:

62. Knoblauch J, Drobnitch ST, Peters WS, Knoblauch M (2016a) In situ microscopy reveals reversible cell wall swelling in kelp sieve tubes: one mechanism for turgor generation and flow control? Plant Cell Environ 39:1727–1736. 10.1111/pce.12736

63. Knoblauch J, Peters WS, Knoblauch M (2016b) The gelatinous extracellular matrix facilitates transport studies in kelp: visualization of pressure-induced flow reversal across sieve plates. Ann Bot 117:599–606. 10.1093/aob/mcw007

64. Koehl M, Alberte RS (1988) Flow, flapping, and photosynthesis of *Nereocystis luetkeana*: a functional comparison of undulate and flat blade morphologies. Mar Biol 99:435–444. 10.1007/BF02112137

65. Koehl M, Silk WK, Liang H, Mahadevan L (2008) How kelp produce blade shapes suited to different flow regimes: A new wrinkle. Integr Comp Biol 48:834–851. 10.1093/icb/icn069

66. Koehl M, Wainwright SA (1977) Mechanical adaptations of a giant kelp. Limnol Oceanogr 22:1067–1071. 10.4319/lo.1977.22.6.1067

67. Konar B, Edwards MS, Bland A, et al (2017) A swath across the great divide: Kelp forests across the Samalga Pass biogeographic break. Cont Shelf Res 143:78–88. 10.1016/j.csr.2017.06.007

68. Kuo A, Bushnell B, Grigoriev IV (2014) Chapter One – Fungal Genomics: Sequencing and Annotation. In: Martin FM (ed) Advances in Botanical Research. Academic Press, pp 1–52

69. Lee S, Bakker CR, Vitzthum C, et al (2022) Pairs and Pairix: a file format and a tool for efficient storage and retrieval for Hi-C read pairs. Bioinforma Oxf Engl 38:1729–1731. 10.1093/bioinformatics/btab870

70. Li H (2013) Aligning sequence reads, clone sequences and assembly contigs with BWA-MEM

132. Liggan LM, Martone PT (2020) Gas composition of developing pneumatocysts in bull kelp *Nereocystis luetkeana* (Phaeophyceae). J Phycol 56:1367–1372. 10.1111/jpy.13037

71. Lind AC, Konar B (2017) Effects of abiotic stressors on kelp early life-history stages. ALGAE 32:223–233. 10.4490/algae.2017.32.8.7

72. Luikart G, Kardos M, Hand B, et al (2018) Population Genomics: Advancing Understanding of Nature. In: Population Genomics:Concepts, Approaches and Applications, Rajora, O. Springer, Cham, pp 3–79

73. Lüning K, Freshwater W (1988) Temperature tolerance of northeast pacific marine algae. J Phycol 24:310–315. 10.1111/j.1529-8817.1988.tb04471.x

74. Lynch M, Trickovic B, Kempes CP (2022) Evolutionary scaling of maximum growth rate with organism size. Sci Rep 12:22586. 10.1038/s41598-022-23626-7

75. Manni M, Berkeley MR, Seppey M, et al (2021) BUSCO Update: Novel and Streamlined Workflows along with Broader and Deeper Phylogenetic Coverage for Scoring of Eukaryotic, Prokaryotic, and Viral Genomes. Mol Biol Evol 38:4647–4654. 10.1093/molbev/msab199

76. Marks RA, Hotaling S, Frandsen PB, VanBuren R (2021) Representation and participation across 20 years of plant genome sequencing. Nat Plants 7:1571–1578. 10.1038/s41477-021-01031-8

77. Maxell B, Miller K (1996) Demographic studies of the annual kelps *Nereocystis luetkeana* and *Costaria costata* (Laminariales, Phaeophyta) in Puget Sound, Washington. Bot Mar 39:479–489. 10.1515/botm.1996.39.1-6.479

78. McPherson ML, Finger DJI, Houskeeper HF, et al (2021) Large-scale shift in the structure of a kelp forest ecosystem co-occurs with an epizootic and marine heatwave. Commun Biol 4:1–9. 10.1038/s42003-021-01827-6

79. Melén K, Krogh A, von Heijne G (2003) Reliability Measures for Membrane Protein Topology Prediction Algorithms. J Mol Biol 327:735–744. 10.1016/S0022-2836(03)00182-7

80. Miller K, Estes J (1989) Western range extension for *Nereocystis luetkeana* in the north pacific ocean. Bot Mar 32:535–538. 10.1515/botm.1989.32.6.535

81. Nagasato C, Motomura T (2009) Effect of Latrunculin B and Brefeldin a on Cytokinesis in the Brown Alga Scytosiphon Lomentaria (scytosiphonales, Phaeophyceae). J Phycol 45:404–412. 10.1111/j.1529-8817.2009.00655.x

82. Nakamura-Gouvea N, Alves-Lima C, Benites LF, et al (2022) Insights into agar and secondary metabolite pathways from the genome of the red alga *Gracilaria domingensis* (Rhodophyta, Gracilariales). J Phycol 58:406–423. 10.1111/jpy.13238

83. Nicholson NL (1970) Field studies on the giant kelp *Nereocystis*. J Phycol 6:177–182. 10.1111/j.1529-8817.1970.tb02378.x

84. Open2C, Abdennur N, Fudenberg G, et al (2023) Pairtools: from sequencing data to chromosome contacts. BioRxiv Prepr Serv Biol 2023.02.13.528389. 10.1101/2023.02.13.528389

85. Pace D (1981) Kelp community development in Barkley Sound, British Columbia following sea urchin removal. In: Proceedings of the International Seaweed Symposium. International Seaweed Association, Bangor, North Wales, pp 457–463

86. Parker BC, Bleck J (1965) A new species of Elk Kelp. Trans San Diego Soc Nat Hist 14:57–64

87. Pfister CA, Berry HD, Mumford T (2018) The dynamics of Kelp Forests in the Northeast Pacific Ocean and the relationship with environmental drivers. J Ecol 106:1520–1533. 10.1111/1365-2745.12908

88. Popovic R, Colbow K, Vidaver W, Bruce D (1983) Evolution of O₂ in Brown Algal Chloroplasts. Plant Physiol 73:889–892

89. Poulson ME, McNeil AJ, Donahue RA (2011) Photosynthetic response of *Nereocystis luetkeana* (Phaeophyta) to high light. Phycol Res 59:156–165. 10.1111/j.1440-1835.2011.00614.x

90. Price AL, Jones NC, Pevzner PA (2005) De novo identification of repeat families in large genomes. Bioinforma Oxf Engl 21 Suppl 1:i351–358. 10.1093/bioinformatics/bti1018

91. Quevillon E, Silventoinen V, Pillai S, et al (2005) InterProScan: protein domains identifier. Nucleic Acids Res 33:W116–120. 10.1093/nar/gki442

92. R Core Team (2024) R: A Language and Environment for Statistical Computing

93. Ramírez F, Bhardwaj V, Arrigoni L, et al (2018) High-resolution TADs reveal DNA sequences underlying genome organization in flies. Nat Commun 9:189. 10.1038/s41467-017-02525-w

94. Ranallo-Benavidez TR, Jaron KS, Schatz MC (2020) GenomeScope 2.0 and Smudgeplot for reference-free profiling of polyploid genomes. Nat Commun 11:1432. 10.1038/s41467-020-14998-3

95. Redmond S, Green L, Yarish C, et al (2014) New England seaweed culture handbook. Connecticut Sea Grant CTSG-14-01. 92

96. Rhie A, McCarthy SA, Fedrigo O, et al (2021) Towards complete and error-free genome assemblies of all vertebrate species. Nature 592:737–746. 10.1038/s41586-021-03451-0

97. Rhie A, Walenz BP, Koren S, Phillippy AM (2020) Merqury: reference-free quality, completeness, and phasing assessment for genome assemblies. Genome Biol 21:245. 10.1186/s13059-020-02134-9

98. Rogers-Bennett L, Catton CA (2019) Marine heat wave and multiple stressors tip bull kelp forest to sea urchin barrens. Sci Rep 9:15050. 10.1038/s41598-019-51114-y

99. Saier MH Jr, Reddy VS, Moreno-Hagelsieb G, et al (2021) The Transporter Classification Database (TCDB): 2021 update. Nucleic Acids Res 49:D461–D467. 10.1093/nar/gkaa1004

100. Salamov AA, Solovyev VV (2000) Ab initio Gene Finding in Drosophila Genomic DNA. Genome Res 10:516–522

101. Schmitz K, Lobban CS (1976) A survey of translocation in laminariales (Phaeophyceae). Mar Biol 36:207–216. 10.1007/BF00389281

102. Schmitz K, Srivastava LM (1976) The Fine Structure of Sieve Elements of *Nereocystis Luetkeana*. Am J Bot 63:679–693. 10.1002/j.1537-2197.1976.tb11856.x

103. Schmitz K, Srivastava LM (1974) The enzymatic incorporation of (32)P into ATP and other organic compounds by sieve-tube sap of Macrocystis integrifolia Bory. Planta 116:85–89. 10.1007/BF00390206

104. Setchell WA (1908) Nereocystis and Pelagophycus. Bot Gaz 45:125–134

105. Setchell WA, Gardner NL (1925) The marine algae of the Pacific coast of North America. University of California Press, Berkeley, Calif

106. Shaffer HB, Toffelmier E, Corbett-Detig RB, et al (2022) Landscape Genomics to Enable Conservation Actions: The California Conservation Genomics Project. J Hered 113:577–588. 10.1093/jhered/esac020

107. Shan T, Yuan J, Su L, et al (2020) First Genome of the Brown Alga *Undaria pinnatifida*: Chromosome-Level Assembly Using PacBio and Hi-C Technologies. Front Genet 11:

108. Sim SB, Corpuz RL, Simmonds TJ, Geib SM (2022) HiFiAdapterFilt, a memory efficient read processing pipeline, prevents occurrence of adapter sequence in PacBio HiFi reads and their negative impacts on genome assembly. BMC Genomics 23:157. 10.1186/s12864-022-08375-1

109. Smale DA (2020) Impacts of ocean warming on kelp forest ecosystems. New Phytol 225:1447–1454. 10.1111/NPH.16107

110. Smit A, Hubley R, Green p (1996) RepeatMasker Open-4.0.

111. Starko S, Mansfield Shawn D., and Martone PT (2018) Cell wall chemistry and tissue structure underlie shifts in material properties of a perennial kelp. Eur J Phycol 53:307–317. 10.1080/09670262.2018.1449013

112. Steele TS, Brunson JK, Maeno Y, et al (2022) Domoic acid biosynthesis in the red alga Chondria armata suggests a complex evolutionary history for toxin production. Proc Natl Acad Sci 119:e2117407119. 10.1073/pnas.2117407119

113. Supratya VP, Coleman LJM, Martone PT (2020) Elevated temperature affects phenotypic plasticity in the bull kelp (*Nereocystis luetkeana*, Phaeophyceae). J Phycol 56:1534–1541. 10.1111/jpy.13049

114. Ter-Hovhannisyan V, Lomsadze A, Chernoff YO, Borodovsky M (2008) Gene prediction in novel fungal genomes using an ab initio algorithm with unsupervised training. Genome Res 18:1979–1990. 10.1101/gr.081612.108

115. Teufel F, Almagro Armenteros JJ, Johansen AR, et al (2022) SignalP 6.0 predicts all five types of signal peptides using protein language models. Nat Biotechnol 40:1023–1025. 10.1038/s41587-021-01156-3

116. Thapa HR, Lin Z, Yi D, et al (2020) Genetic and biochemical reconstitution of bromoform biosynthesis in *Asparagopsis* lends insights into seaweed ROS enzymology. ACS Chem Biol 15:1662–1670. 10.1021/acschembio.0c00299

117. Theissinger K, Fernandes C, Formenti G, et al (2023) How genomics can help biodiversity conservation. Trends Genet 39:545–559. 10.1016/j.tig.2023.01.005

118. Tillich M, Lehwark P, Pellizzer T, et al (2017) GeSeq – versatile and accurate annotation of organelle genomes. Nucleic Acids Res 45:W6–W11. 10.1093/nar/gkx391

119. Turner N, Haga KL, Hulbert AJ, Else PL (2005) Relationship between body size, Na+-K+-ATPase activity, and membrane lipid composition in mammal and bird kidney. Am J Physiol Regul Integr Comp Physiol 288:R301–310. 10.1152/ajpregu.00297.2004

120. Uliano-Silva M, Ferreira JGRN, Krasheninnikova K, et al (2023) MitoHiFi: a python pipeline for mitochondrial genome assembly from PacBio high fidelity reads. BMC Bioinformatics 24:288. 10.1186/s12859-023-05385-y

121. Wade R, Augyte S, Harden M, et al (2020) Macroalgal germplasm banking for conservation, food security, and industry. PLOS Biol 18:e3000641. 10.1371/JOURNAL.PBIO.3000641

122. Walker DC (1980) Sorus abscission from laminae of Nereocystis luetkeana (Mert.) Post. and Rup. University of British Columbia

123. Wang X, Shao Z, Fu W, et al (2013) Chloroplast genome of one brown seaweed, *Saccharina japonica* (Laminariales, Phaeophyta): Its structural features and phylogenetic analyses with other photosynthetic plastids. Mar Genomics 10:1–9. 10.1016/j.margen.2012.12.002

124. Wernberg T, Coleman MA, Bennett S, et al (2018) Genetic diversity and kelp forest vulnerability to climatic stress. Sci Rep 8:. 10.1038/s41598-018-20009-9

125. Wheeler WN, Smith RG, Srivastava LM (1984) Seasonal photosynthetic performance of *Nereocystis luetkeana*. Can J Bot 62:664–670. 10.1139/b84-099

126. Wickham H (2009) ggplot2: Elegant Graphics for Data Analysis. Springer, New York, NY

127. Wilson MD, Riemer C, Martindale DW, et al (2001) Comparative analysis of the gene-dense ACHE/TFR2 region on human chromosome 7q22 with the orthologous region on mouse chromosome 5. Nucleic Acids Res 29:1352–1365. 10.1093/nar/29.6.1352

128. Ye N, Zhang X, Miao M, et al (2015) Saccharina genomes provide novel insight into kelp biology. Nat Commun 6:6986. 10.1038/ncomms7986

129. Zhang M, Wang Y, Chen X, et al (2021) Plasma membrane H+-ATPase overexpression increases rice yield via simultaneous enhancement of nutrient uptake and photosynthesis. Nat Commun 12:735. 10.1038/s41467-021-20964-4

130. Zheng Z, Chen H, Wang H, et al (2019) Characterization of the complete mitochondrial genome of bull kelp, Nereocystis luetkeana. Mitochondrial DNA Part B 4:630–631. 10.1080/23802359.2018.1501295

131. Zhou K, Salamov A, Kuo A, et al (2015) Alternative splicing acting as a bridge in evolution. Stem Cell Investig 2:19. 10.3978/j.issn.2306-9759.2015.10.01

