## Supplementary table 1 for "The reference genome for the northeastern Pacific bull kelp, *Nereocystis luetkeana*"

Suplementary table 1: Contamination screening and taxonomy assignment of non Laminariales contigs in *Nereocystis luetkeana* genome

| | # contigs | Cumulative length |
|---|---|---|
| **Eukaryota-undef** | 10 | 6,686,382 |
| **Bacteria** | 489 | 100,930,254 |
| **Metazoa** | 14 | 6,431,548 |
| **Viridiplantae** | 8 | 3,361,602 |
| **Virus** | 1 | 56,540 |
| **Undef** | 2 | 217,266 |
| **Bamfordvirae** | 1 | 1,254,679 |
| **Total** | 525 | 118,938,271 |
